# Conformational landscape of soluble α-klotho revealed by cryogenic electron microscopy

**DOI:** 10.1101/2024.03.02.583144

**Authors:** Nicholas J. Schnicker, Zhen Xu, Mohammad Amir, Lokesh Gakhar, Chou-Long Huang

**Affiliations:** Protein and Crystallography Facility, University of Iowa Carver College of Medicine, Iowa City, Iowa, 52242, USA; Department of Molecular Physiology and Biophysics, University of Iowa Carver College of Medicine, Iowa City, Iowa, 52242, USA; Department of Internal Medicine, University of Iowa Carver College of Medicine, Iowa City, Iowa, 52242, USA; Department of Biochemistry and Molecular Biology, University of Iowa, Iowa City, Iowa, 52242, USA

**Author notes:** Equal contribution (Nicholas J. Schnicker, Zhen Xu, Mohammad Amir).

**Keywords:** Soluble alpha-klotho (sKLA), Fibroblast growth factor (FGF), cryogenic electron microscopy (cryo-EM), ternary complex, monomer, dimer, 3D variability analysis (3DVA)

## Abstract

α-Klotho (KLA) is a type-1 membranous protein that can associate with fibroblast growth factor receptor (FGFR) to form co-receptor for FGF23. The ectodomain of unassociated KLA is shed as soluble KLA (sKLA) to exert FGFR/FGF23-independent pleiotropic functions. The previously determined X-ray crystal structure of the extracellular region of sKLA in complex with FGF23 and FGFR1c suggests that sKLA functions solely as an on-demand coreceptor for FGF23. To understand the FGFR/FGF23-independent pleiotropic functions of sKLA, we investigated biophysical properties and structure of apo-sKLA. Mass photometry revealed that sKLA can form a stable structure with FGFR and/or FGF23 as well as sKLA dimer in solution. Single particle cryogenic electron microscopy (cryo-EM) supported the dimeric structure of sKLA. Cryo-EM further revealed a 3.3Å resolution structure of apo-sKLA that overlays well with its counterpart in the ternary complex with several distinct features. Compared to the ternary complex, the KL2 domain of apo-sKLA is more flexible. 3D variability analysis revealed that apo-sKLA adopts conformations with different KL1-KL2 interdomain bending and rotational angles. The potential multiple forms and shapes of sKLA support its role as FGFR-independent hormone with pleiotropic functions. A comprehensive understanding of the sKLA conformational landscape will provide the foundation for developing klotho-related therapies for diseases.

## INTRODUCTION

α-Klotho (KLA) is a type I single-pass transmembrane protein consisting of 1012 amino acids (human Klotho) with a large extracellular region (1). The ectodomain contains two homologous repeats named KL1 and KL2, which are comprised from residues 57-506 and 515-953, respectively, and has multiple N- and O-linked glycosylation sites. The ectodomain is followed by the transmembrane-spanning segment and a short 11 amino acids intracellular carboxyl terminus. KLA is abundantly produced in the kidney and several regions in the brain and exerts anti-aging effects (2–4). Mice homozygous for a hypomorphic *klotho* allele (*kl*/*kl*) die prematurely at around 2-3 months of age. The full-length membranous KLA can associate with fibroblast growth factor receptors (FGFR) to form co-receptors for the ligand fibroblast growth factor-23 (FGF23). FGF23 is a bone-derived circulating hormone that is important in calcium and phosphate metabolism (5–7). KLA-deficient mice have severe hyperphosphatemia due to defects in the KLA-FGF23-vitamin D regulatory axis (8–10). Phosphate retention is pivotal for growth retardation and premature death of klotho-deficient mice; dietary phosphate restriction rescues growth defects and premature death of the mice (8–10). Of note, the affinity between KLA and FGFR is relatively low; FGF23 binding to the coreceptor facilitates the interaction and formation of the ternary FGF23-FGFR-KLA complexes (11). Without FGF23, significant fractions of membranous KLA are dissociated from FGFR existing in a free form (11). The ectodomain of free KLA can be cleaved by metalloproteases (ADAM10/17) and released as soluble KLA (sKLA) into the systemic circulation, urine, and cerebrospinal fluid (12).

The function of Klotho family proteins to form coreceptors for FGFs and determine the specificity for FGFs are well established (13–15). KLA associates with FGFR1c, FGFR3c, and FGFR4 to form specific coreceptor complexes with FGF23, whereas β-Klotho (KLB) associates with FGFR1c and FGFR4 to form complexes with FGF21 and FGF19, respectively. Despite the overall similar domain structure to KLA, the ectodomain of KLB is not known to be cleaved. Many studies have demonstrated that the cleaved sKLA can act independently of FGFR coreceptor as an endocrine/local paracrine hormone with pleiotropic effects (4,16–18). For example, sKLA regulates the activity of ion channels and transporters, inhibits Wnt and TGF-β1 signaling to suppress cellular senescence, tissue fibrosis, and cancer metastasis. sKLA also exerts inhibition on insulin and growth factor driven PI3K/Akt signaling contributing to extension of lifespan in mice, cardioprotection, and inhibition of tumor cell proliferation.

An X-ray crystal structure for sKLA in ternary complex with a truncated FGF23 and the FGF23-binding domains (D2 and D3 domain) of FGFR1c has been solved (PDB ID 5W21) (15). A large receptor binding arm (RBA, residues 530-578) of the KL2 domain engages interactions with D2 and D3 of FGFR1c. The KL1 and KL2 domains are connected by a proline-rich linker (residues 507-514) and further stabilized by a zinc coordination site comprised of residues D426, C739, D745, and D815 (15). The N-terminus of FGF23 is bound between the D2 and D3 domains of FGFR, and the truncated C-terminus of FGF23 interacts with sKLA at the inter KL1-KL2 cleft and extends into the KL2 domain. It is known that both the KL1 and KL2 domains of KLA share sequence and structural homology to family 1 glycosidases (1). The KL1 and KL2 domain of KLA each contains a cavity equivalent to the catalytic center of the family 1 glycosidases. Interestingly, while both domains lack an essential catalytic glutamate residue typically present in the family 1 glycosidase active sites, glycosidase activity has been observed for KLA by multiple groups (19–21). The ternary complex X-ray structure shows that the access to the active site is blocked by a β6α6 loop. The lack of a catalytic glutamate residue and blocked access to the active site in the ternary sKLA structure have raised questions as to whether sKLA functions (such as enzymatic glycosidase activity) independently of FGF23 and FGFR.

To better comprehend the pleiotropic FGF23-independent functions of sKLA, the present study investigates the biophysical properties and structure of sKLA in the absence of binding partners using size exclusion chromatography (SEC), mass photometry (MP), and single particle cryogenic electron microscopy (cryo-EM). We found that apo-sKLA can form a pseudodimer and monomeric sKLA adopts multiple conformations. The results support the notion that sKLA could have FGF23-independent functions as well as regulating FGF23 signaling.

## RESULTS

### Biophysical characterization of sKLA

To investigate potential pleiotropic functions of sKLA, a commercial source of recombinant sKLA was characterized using size-exclusion chromatography (SEC) and mass photometry (MP). MP is a single molecule-based technique to determine the mass distribution of macromolecules in solution. The SEC chromatograph showed the presence of two species of sKLA consistent with dimeric structure (elution volume 11.1 mL, ∼364 kDa based on molecular weight standards) in addition to monomeric sKLA (12.9 mL, ∼155 kDa) (Figure 1A). The commercial source of sKLA is provided in the absence of reducing agents. To determine if the formation of the dimeric sKLA was redox-dependent, the peak fractions were pooled, concentrated, and reinjected onto the SEC column either with or without reducing agent (DTT) incubation. Dimeric sKLA was reduced almost completely to the monomeric form in the presence of DTT (Figure 1B). Some variability in the amount of the dimer was observed from various lots of sKLA supplied by R&D Systems (data not shown). Confirmation of molecular weight estimates from SEC were performed by collecting MP data on monomeric and dimeric peak fractions from SEC. Representative MP data shows a dimeric (Figure 1C) and monomeric (Figure 1D) species from corresponding SEC fractions. The results provide evidence of the existence of a dimeric form of sKLA, which has previously been reported (22). The nature of dimer formation remains unknown as attempts to reform the dimer from the monomeric species under oxidizing buffer conditions were unsuccessful (data not shown).

**Figure 1.**
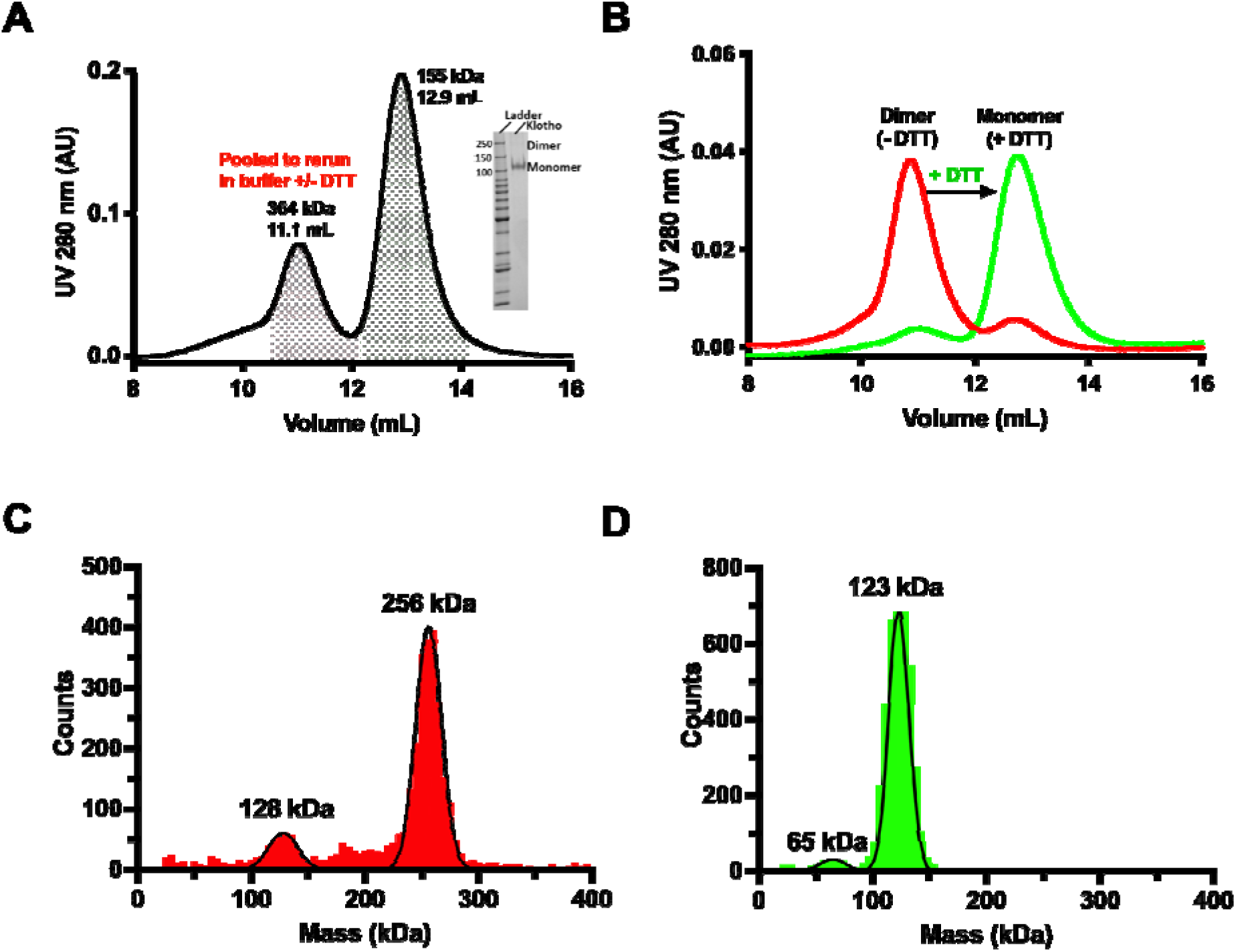
sKLA size-exclusion profiles and mass photometry measurements. **(A)** A representative size-exclusion chromatogram (SEC) of sKLA depicting dimeric form (molecular weight, 364 kDa) at ∼11 mL and monomeric sKLA (molecular weight, 155 kDa) at 13 mL based on standard molecular weight. SDS-PAGE (inset) shows the monomer and dimer sKLA. **(B)** Dimeric sKlotho from panel A was pooled and incubated with DTT and rerun onto SEC. **(C)** Mass photometry analysis of dimeric sKLA depicts enrichment of dimeric form (63%) compared to monomeric form (11%). **(D)** Mass photometry of monomeric sKLA showing enrichment of monomer sKLA (90%) as well as cleavable KL1/KL2 domain of sKLA.

Next, MP was used to examine whether sKLA can form independent complexes with FGF23 and/or FGFR1c. In a solution containing 10 nM sKLA and 10 nM FGF23, 18% of sKLA formed a complex with FGF23 with the rest remaining as free sKLA (Figure 2A). Free FGF23 is smaller than the 30 kDa lower limit of detection for MP and thus is not observed. In a solution containing 10 nM sKLA and 10 nM FGFR1c, 35% of sKLA formed a complex with FGFR1c with the rest remaining as free sKLA (Figure 2B). In a mixture of all three proteins with a 1:4:4 molar ratio of sKLA: FGFR1c: FGF23, almost all variations of free and bound species can be observed (Figure 2C). Distinct peaks for free sKLA and sKLA complexes with FGF23 (20%), FGFR1c (29%), and both FGF23 and FGFR1c (28%) are present (Figure 2D). No binding between FGF23 and FGFR1c was detected. The results support the notion that the affinity of FGF23 for FGFR is weak and KLA increases the affinity of KLA-FGFR for FGF23.

**Figure 2.**
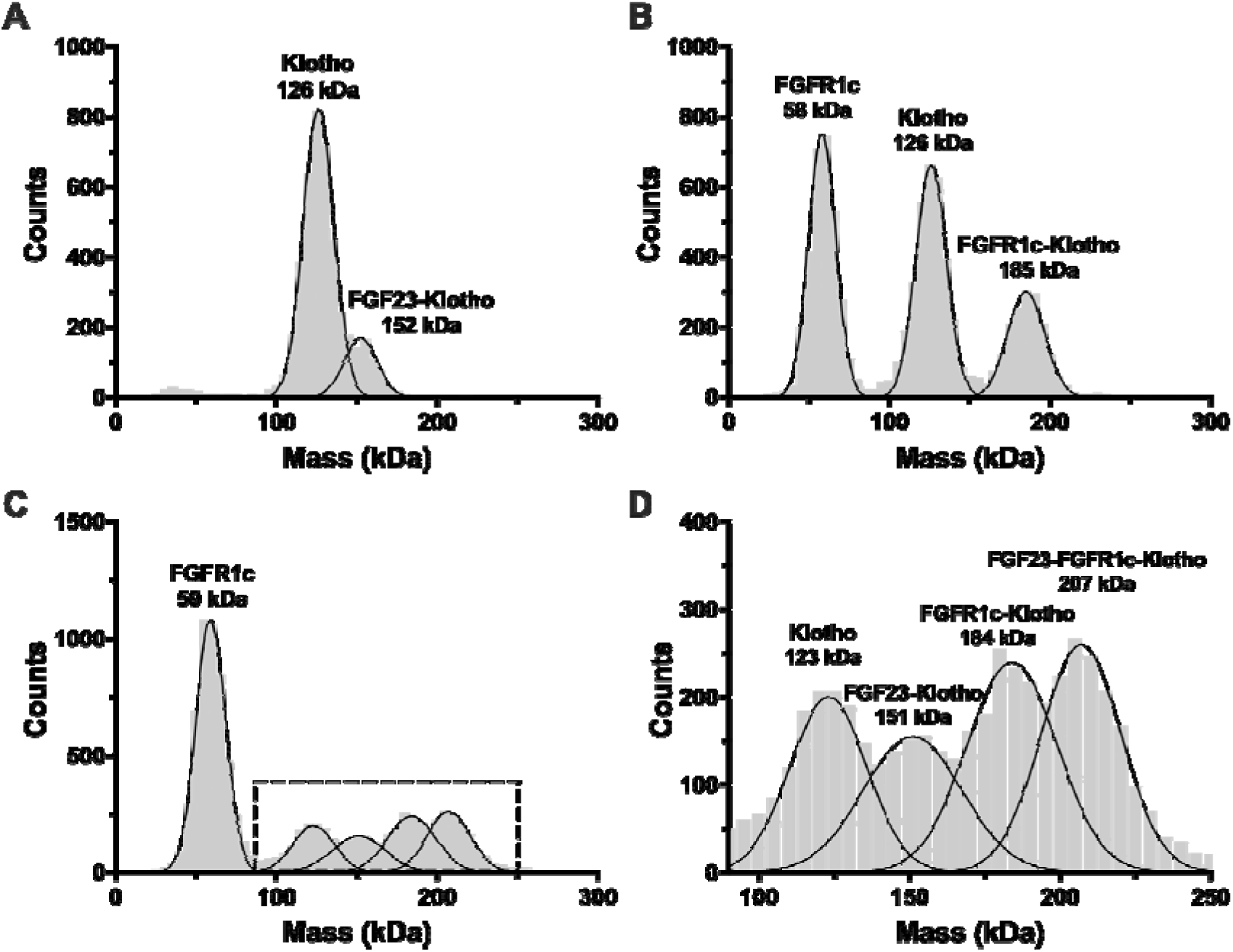
Mass photometry analysis of complexes formation of sKLA with FGFR1c and FGF23. **(A)** Mass distribution of 10 nM sKLA with FGF23 in 1:1 molar ratio. **(B)** Mass distribution of 10 nM sKLA with FGFR1c in 1:1 molar ratio. **(C)** Mass distribution of 10 nM sKLA with FGFR1c and FGF23 in 1:4:4 molar ratio. **(D)** Zoom-in view of region in panel C. The peaks in mass distribution are associated with FGFR1c (59 kDa), sKLA (123 kDa), sKLA-FGF23 1:1 complex (151 kDa), sKLA-FGFR1c 1:1 complex (184 kDa) and sKLA-FGFR1c-FGF23 1:1:1 complex (207 kDa).

### Cryo-EM structure of sKLA monomer

The single particle cryo-EM reconstruction workflow for the sKLA monomer (Table 1, Figure 3A) yielded a 3D reconstruction with an overall gold-standard Fourier shell correlation (FSC) resolution of 3.3 Å (Figure 3B). The workflow included template-based particle picking (from 2D class averages), 2D class average rebalancing (to remove particles with oversampled views), and 3D classification. The local resolution ranges from 3.0 Å in the inner regions of the glycoside hydrolase family-1 (GH1) domains of KL1 and KL2 to ∼5.0 Å near the RBA loop surface region in KL2 (Figure S1). The structure has dynamic regions that are too disordered and therefore don’t have visible density, such as the β1α1 loop region from residues 98-119 and the RBA (Figure 3C). The metal binding site between residues D426, C739, D745, and D815 is not well resolved, and therefore does not contain a metal modeled in the site (see arrow in Figure 3C). In the sKLA ternary complex crystal structure, this site was occupied by zinc. Apo sKLA overlays well onto the sKLA-FGF23-FGFR1c ternary structure with an overall backbone RMSD of 1.54 Å (Figure 3D). The individual domains align better with a backbone RMSD of 0.98 and 1.0 Å for KL1 and KL2, respectively.

**Figure 3.**
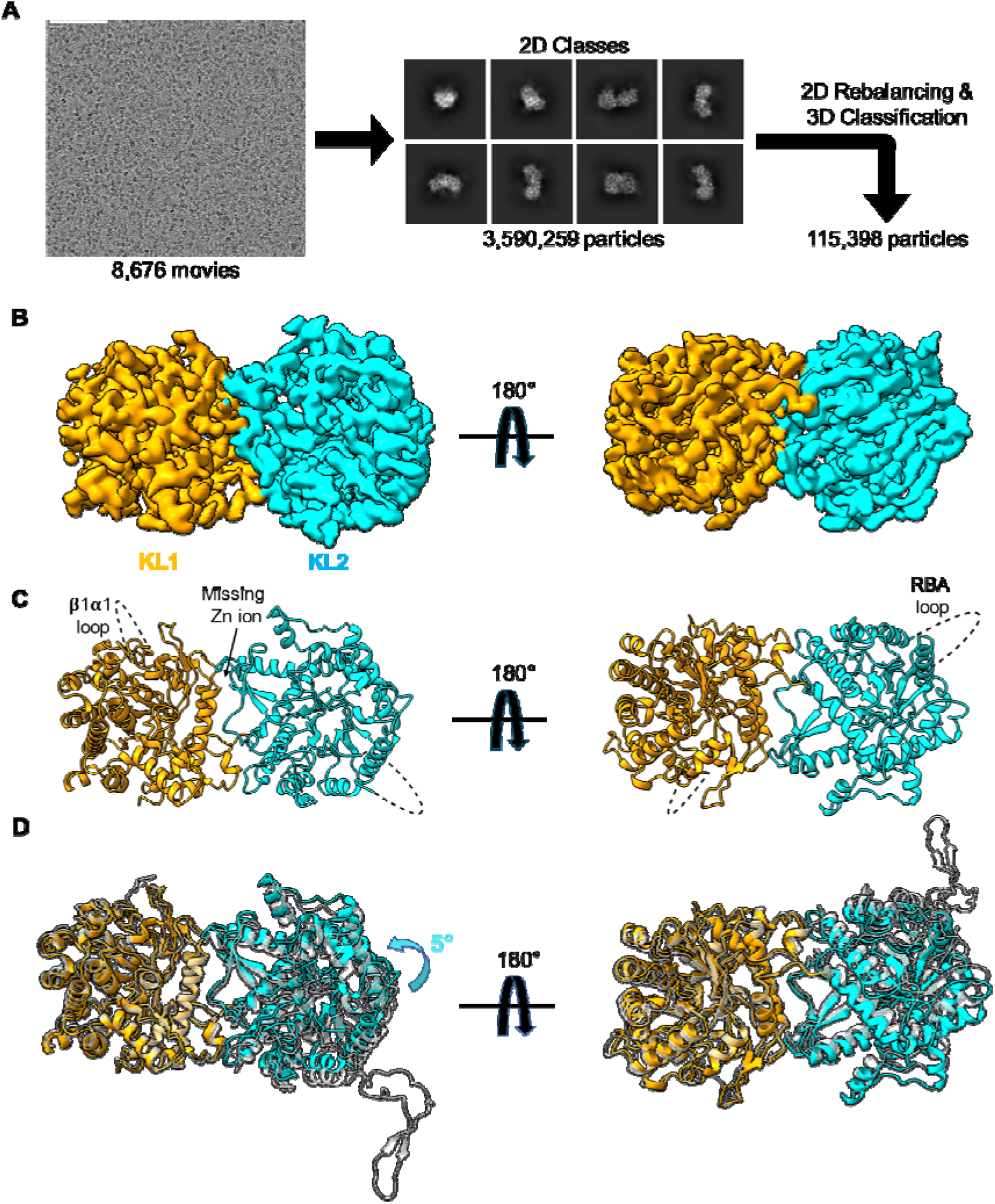
Overview of sKLA monomer cryo-EM analysis and 3D reconstruction with final model. **(A)** A representative electron micrograph of sKLA at defocus range of -1.0 to -2.5 μm (left), and representative 2D class averages (right). **(B)** 3D reconstruction using non-uniform refinement for monomeric sKLA at 3.3LÅ resolution from 115,398 particles. **(C)** Final structure of monomeric sKLA showing missing residue regions. **(D)** Comparison with sKLA X-ray structure (PDB: 5W21, gray) aligned on the KL1 domain. Note that the density of RBA is invisible in apo-sKLA cryo-EM structure in contrast to the ternary complex structure with FGFR and FGF23.

**Table 1.** Cryo-EM data collection and model refinement statistics.

|  | sKLA monomer | sKLA dimer |
| --- | --- | --- |
| ***Data collection and image processing*** |  |  |
| Microscope | Titan Krios | Titan Krios |
| Voltage (kV) | 300 | 300 |
| Camera | K3 | K3 |
| Magnification | 130,000 x | 130,000 x |
| Electron exposure (e<sup>-</sup>/Å<sup>2</sup>) | 50 | 60 |
| Exposure time (s) | 1.099 | 2 |
| Number of frames | 50 | 50 |
| Defocus range (mm) | -1.0 to -2.5 | -0.8 to -2.5 |
| Super resolution pixel size (Å) | 0.32535 | 0.265 |
| Number of movies | 8,676 | 5,760 |
| Particles for final reconstruction (no.) | 115,398 | 30,219 |
| Map resolution (Å) | 3.3 | 6.5 |
| Half map FSC threshold | 0.143 | 0.143 |
| EMDB accession code | 41452 | 42186 |
| ***Model refinement statistics*** |  |  |
| ***Model composition*** |  |  |
| Non-hydrogen atoms | 7,022 | 14,050 |
| Protein residues | 860 | 1720 |
| Ligands | 0 | 0 |
| ***Cross correlation*** |  |  |
| Mask | 0.65 | 0.53 |
| Volume | 0.65 | 0.52 |
| ***RMSD*** |  |  |
| Bond length (Å) | 0.004 | 0.004 |
| Bond Angles (°) | 0.969 | 1.042 |
| ***Ramachandran*** |  |  |
| Favored (%) | 92.74 | 89.46 |
| Allowed (%) | 7.26 | 10.54 |
| Outlier (%) | 0.00 | 0.00 |
| ***Validation*** |  |  |
| MolProbity score | 2.11 | 2.26 |
| Clashscore | 13.84 | 15.51 |
| Rotamer outliers (%) | 0.00 | 0.95 |
| PDB accession code | 8TOH | 8UF8 |

### Pseudodimer structure of sKLA

Studies of sKLA have predominantly focused on the idea that sKLA exists in the monomeric form (∼125 kDa). Yet, several studies have reported the existence of a dimeric form of sKLA that could be functionally distinctive from the FGF23 co-receptor function (23,24). Our dimeric sKLA cryo-EM structure was determined from a ∼6.5 Å reconstruction (Figure 4A). As the number of dimer particles were limited, a higher resolution structure for the dimer was not obtained. Instead, the sKLA monomer structure was fit into the dimeric map to observe the overall organization. The interface is formed at two nearby regions through the KL2 domains of both protomers. The total dimer interface is small (∼400 Å^2^). Residues involved in the sKLA dimer formation are shown in Figure 4B. Analysis using PDBePISA (25) revealed no obvious hydrogen bond nor electrostatic based interactions at the low-resolution interface. Weak dimeric interactions are often formed through hydrophobic interactions (26). Due to the small interface and lack of strong interactions at the interface, we refer to this form as a ‘pseudodimeric’ arrangement. To test whether the pseudodimeric arrangement could be feasible in both soluble and membrane bound forms, the full-length membranous KLA model from the AlphaFold Structural Database (https://alphafold.ebi.ac.uk/) was overlaid onto each protomer of the pseudodimer and was observed to fit well without any clashes (Figure 4C). Notably, recent cryo-EM structures solved by Chen et al. of sKLA in complex with FGF23 and several FGFRs also observed dimeric sKLA molecules (Figure S2A, S2B) (22). The density for the second sKLA molecule in the Chen et al. structures wasn’t as strong, and therefore was not utilized while building the main structural models. An overlay of one of the maps from Chen et al. with the sKLA dimer map from this study shows very good agreement of the arrangement and interface (Figure S2C), supporting the notion that the sKLA dimer could exist in multiple scenarios. While checking for the possibility of intermolecular disulfides, no Cys residues were found at the dimer interface. The distribution of Cys residues in relation to the dimer interface is shown in Figure S3.

**Figure 4.**
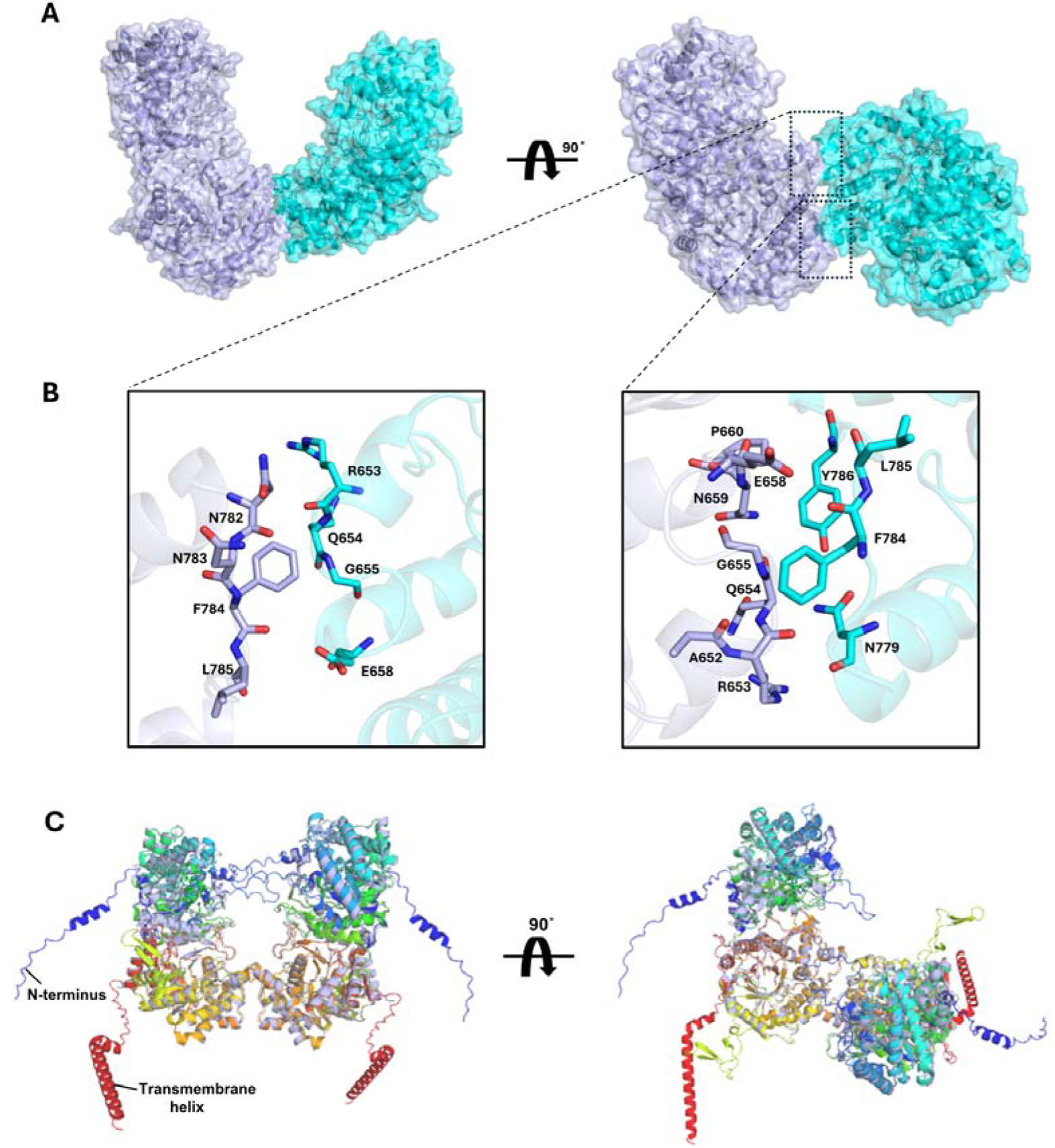
sKLA pseudodimeric structure. **(A)** 3D reconstruction (transparent map) of sKLA dimer with 2 monomers (cartoons) rigid body refined into the EM map. Note contacts at two nearby interface regions. **(B)** Potential residues involved in the dimeric interface. **(C)** The sKLA dimer model overlaid with the sKLA model from the AlphaFold Protein Structure Database. The membrane spanning region of KLA predicted from the AlphaFold Protein Structure Database is colored red.

### Variability analysis of monomeric sKLA

To observe the continuous conformational dynamics of sKLA, 3D variability analysis (3DVA) in cryoSPARC was performed (27). Unlike in standard 3D cryo-EM refinement where a single 3D reconstruction is obtained, 3DVA allows reconstruction of a continuous family of 3D structures to capture discrete and flexible conformations present in the data. In the 3DVA we observed multiple variability components. Among them, we focus on two principal components that display prominent KL1-KL2 interdomain angle bending and rotation movements (Figure 5, Supplementary movie 1 and 2). To determine whether the binding with FGF23 and FGFR may impact sKLA conformation, the sKLA monomer from 3DVA was compared with the sKLA-FGF23-FGFR1c ternary complex crystal structure (5W21). The first variability component displays motion that produces bending between KL1 and KL2 and also a clockwise rotation within KL2 (Figure 5A and B). Comparing the beginning (Component 1 state a, C1a) and end (C1b) states with the 5W21 structure, the KL1 domains align well and therefore have been aligned for the following analyses of 3DVA data. In the 5W21 structure, the relative KL1-KL2 interdomain angle is determined 107° (Figure 5B, left). This angle is determined by defining three points within sKLA, E354 alpha carbon in KL1, zinc atom, and T614 alpha carbon in KL2 (Figure 5A). The most bent state of 3DVA (C1a) aligns well with 5W21 with only 1° angle difference (Figure 5B, middle). The 3DVA bending motion causes an 8° opening between KL1 and KL2 relative to 5W21 structure (Figure 5B, right). The clockwise rotational movement of several KL2 helices for the variability component 1 is highlighted in Figure 5A (see clockwise arrows and Supplementary Movies 1 & 2). The second variability component contains a counter-clockwise twisting motion of KL2 (Figure 5C and D). Figure 5C illustrates that the KL2 helices undergo 6-9° counter-clockwise rotation relative to the sKLA complex crystal structure (see counterclockwise arrows). The angles were determined by using the *angle between helices* script in *PyMOL* (28). Figure 5D illustrates that the least and most counterclockwise rotated states C2a and C2b with 6.7° and 9.3° relative to 5W21, respectively (see Supplementary Movies 3 & 4). Overall, sKLA in the ternary complex overlaid better with the bent or more compact forms from 3DVA (C1a and C2a), suggesting the ternary complex adopts the bent form which allows both KL1 and KL2 to interact with the C-terminus of FGF23. On the other hand, the more open state of 3DVA (C1b and C2b) likely reflects ligand-free sKLA. Analysis of the Debye–Waller (“B”) factors for the consensus sKLA structure suggests that much of the movement is derived from the KL2 domain. The KL1 domain had a lower average B-factor (89 Å^2^) compared to the average KL2 domain B-factor (104 Å^2^, Figure S4).

**Figure 5.**
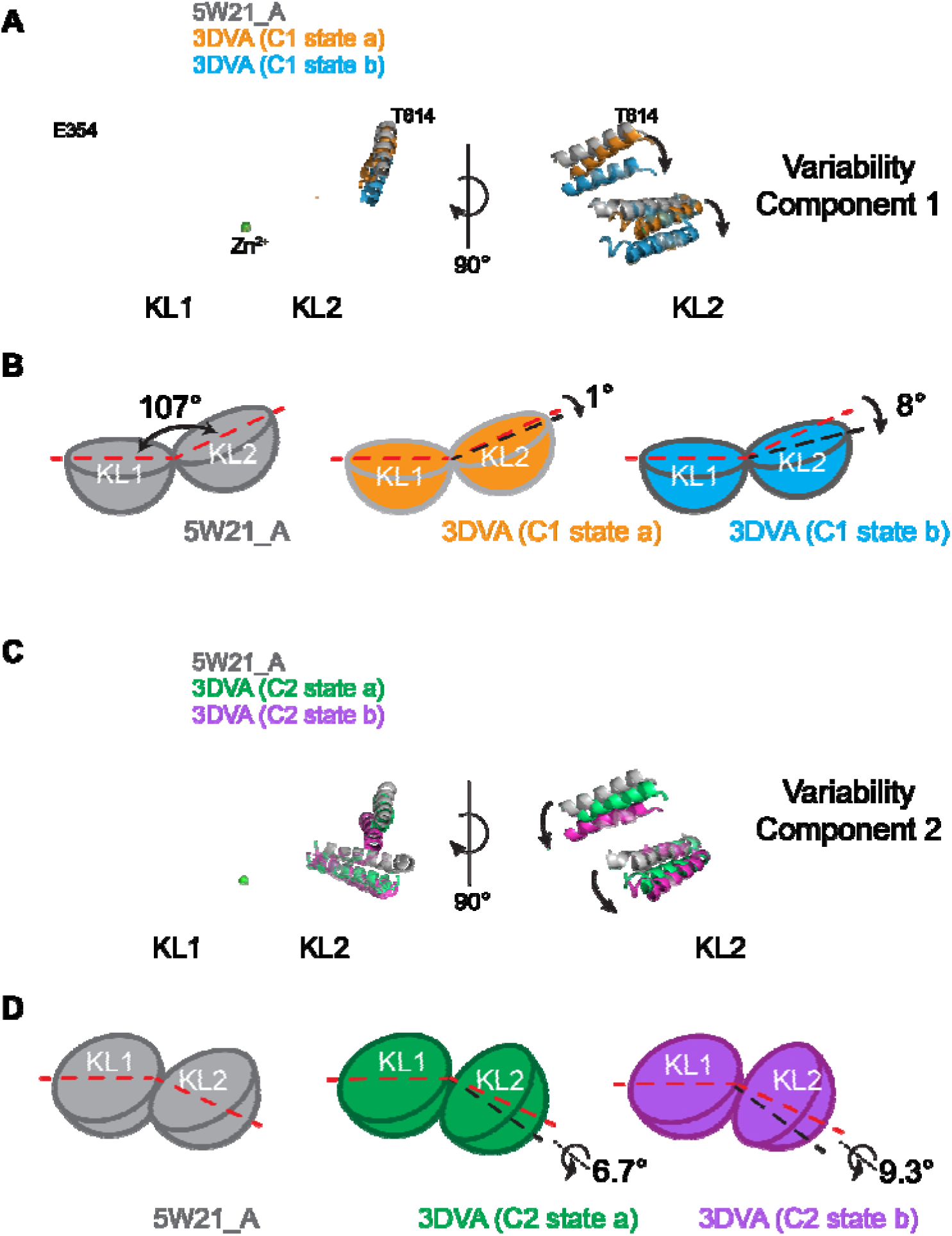
Conformational landscape of sKLA. 3DVA performed on the sKLA monomer cryo-EM data. **(A)** Comparison of the more bent conformation (C1a, orange) and the more open conformation (C1b, blue) from the first variability component. Left, the C1a (orange) and C1b (blue) conformations from 3DVA overlaid with the sKLA ternary complex crystal structure (5W21, gray). The green sphere is a zinc ion from 5W21. Right, a closer view of the 3DVA conformations with the 5W21 structure. Note clockwise rotation indicated by arrows (see Supplementary Movies 1 & 2). **(B)** Cartoon representation of sKLA KL1-KL2 interdomain angles from the first 3DVA component compared with 5W21. **(C)** Comparison of the less rotated conformation (C2a, green) and the more rotated conformation (C2b, purple) from the second variability component. Left and right views are the same as in panel A. Note counterclockwise rotation indicated by arrows (see Supplementary Movies 3 & 4). **(D)** Cartoon representation of sKLA KL1-KL2 interdomain angles from the second 3DVA component compared with 5W21.

## DISCUSSION

Structural studies of KLA have focused on its role in FGF-related signaling mediated by membranous KLA-FGFR-FGF23 complexes (15,22). While functional studies have shown the cleaved sKLA has FGF23-independent pleiotropic functions, the structure of apo-sKLA is not available. Moreover, sKLA either in FGF23-independent state or in the FGF23-bound complexes is generally thought to be monomeric. Some studies have indicated the possible existence of dimeric forms (23,24). Using cryo-EM we reveal the structure of apo-sKLA in FGF23 and FGFR-independent state and a pseudodimer KLA structure.

The structural fold of KL1 and KL2 domains of KLA resembles the glycoside hydrolase family 1 (GH1) enzymes which contain diverse substrates and mechanisms (29,30). The GH1 fold features a triosephosphate isomerase (TIM) barrel fold, comprising a central core formed by β-strands stabilized by α-helices on the surface (31,32). Within the TIM barrel fold are two highly conserved glutamates, one acting as a nucleophile and one as an acid/base. Both KL1 and KL2 domains lack one of the conserved catalytic glutamate residues (31). These residues are N239 and E414 in KL1 and E689 and S872 in KL2. The lack of these conserved glutamates raises the question regarding the enzymatic activity of KLA (15). However, some GH1 enzymes have replaced one of the catalytic glutamate residues with an asparagine or other amino acids and several groups have reported apparently weak glycosidase activity for sKLA (19–21,29,30).

A recent study by Zhong et al. revealed intriguing results on how post-translational modifications on sKLA can affect its FGF23 co-receptor function and glycosidase activity (33). Zhong et al. produced sKLA in both Chinese hamster ovary (CHO) and human embryonic kidney (HEK) 293 cells and found they possess differing glycosylation. CHO-derived sKLA was found to have less FGF23 co-receptor activity compared with transient expression of sKLA in HEK293 cells. Also, CHO-derived sKLA possessed a low-level glycosidase activity whereas HEK sKLA had no measurable activity. A double mutant form of CHO sKLA that introduced the missing glutamate into the KL1 and KL2 domains (N239E and S872E, respectively) was found to have 25-fold higher b-glucuronidase activity compared to the WT protein. Additionally, this double mutant of sKLA had reduced FGF signaling.

One common mode of controlling protein function is through oligomerization. GH1 members have been found to commonly form both dimers and tetramers (34,35). Our results indicate that KLA may also exist in dimers in some settings. While the interface area for the observed sKLA dimer is small, protein interaction interface area can vary widely (36). It is possible that membrane bound KLA or a missing ligand could increase the interface area for the KLA dimer. Interestingly, while reducing agents appear to convert dimers to monomers in SEC and MP experiments, no Cys residue is present at the interface observed in the sKLA pseudodimer cryo-EM structure. The possibility of a non-specific intermolecular disulfide bond resulting in the observed SEC and MP data cannot be excluded, however no Cys on the interface. Alternatively, a redox sensitive intramolecular disulfide may promote specific conformations favorable for dimer formation. In the sKLA-FGF23-FGFR1c ternary complex crystal structure, C572 and C621 form a disulfide and are thought to confer rigidity to the RBA loop (15). A change in the RBA’s rigidity could conceivably play a role in regulating various functions of sKLA. This possibility cannot be assessed in the current apo-sKLA structure because the RBA is too dynamic to observe any density.

Besides KLA and KLB, lactase-phlorizin hydrolase (LPH) is the only GH1 family member that possesses two or more copies of the GH1 domain (34). LPH contains four repeats of the GH1 domain and, looking at the model in the AlphaFold Structural Database, it’s plausible that LPH could adopt a similar overall organization to the sKLA pseudodimer observed in this study (Figure S5, Figure 4C). The implication of KLA dimerization in FGF23-dependent or - independent function remains unknown. It is tempting to speculate that dimerization of sKLA provides complementation for missing nucleophile and acid-base glutamate for KL1 and KL2 and posttranslational modification may influence the propensity for dimer formation. The precise factors for triggering KLA dimerization and possible biological functions of this arrangement will need to be further explored both in its soluble and membrane bound forms.

Without an apo structure for sKLA, it is difficult to assess the sKLA conformational landscape from multi-component complex structures. Crystal contacts present in macromolecular crystal structures can restrict conformational accessibility; therefore cryo-EM was used to solve the monomeric structure of sKLA. The advantage of cryo-EM is that it allows examining of individual macromolecules in solution to uncover multiple conformational states in a more “native” environment. The 3DVA of the sKLA monomer cryo-EM data revealed a more open and a similarly bent conformation as to the sKLA ternary complex crystal structure. Similar conformational changes have been observed with X-ray crystal structures of apo sKLB compared with sKLB bound to C-terminal peptides of FGF19 or FGF21 (13). In sKLB structures bound to FGF19 and FGF21, there is a 17° and 6° closing of the sKLB KL2 domain towards the KL1 domain (interdomain angles), respectively. These differences in the structures of sKLB-FGF complexes are believed to be important for the underlying mechanism of FGF19 and FGF21 specificity and activity. In addition, there is a rotational (twist) movement around the axis parallel to a line joining the centers of mass of KL1 and KL2 domains. In the present study, we compare 3DVA of apo sKLA with the sKLA-FGF23-FGFR1c ternary complex crystal structure (5W21) and find similarities to the observations with sKLB. The sKLA ternary complex overlays better with the more compact, bent conformation (C1a) observed in 3DVA (Figure 5A). The first variability component conformations (C1a and C1b) reflect a 7° difference in the KL1-KL2 interdomain angle (8° relative to 5W21 for open conformation). The ability to adopt the more bent (C1a) state allows sKLA to make interactions with FGF23 from both KL1 and KL2. The more open (C1b) state could serve to regulate FGF signaling or promote alternative functions. The second variability component displayed motion that could also help to regulate FGF signaling as several helices in KL2 undergo a ∼9° rotation (relative to 5W21) near the RBA (Figure 5C). The movement of these helices could destabilize the interaction with the FGFRs, which rely on extensive interactions with the RBA. In the case of sKLB, the changes in interdomain angles for sKLB could be artefacts influenced by bound nanobodies or crystal packing. The fact that our findings for sKLA using cryo-EM are similar to the results from Kuzina et al. (28) support the notion that the conformational landscape of sKLA or sKLB can be complex either in the absence or presence of ligands.

In the ternary complex structure of sKLA-FGFR-FGF23, the access to the GH1 cavity present in KL1 of sKLA is blocked by the β6α6 loop (15). This observation has led Chen et al. to suggest that sKLA does not exert FGF23-independent activity and solely functions as an on-demand FGFR coreceptor for FGF23 signaling. However, weak glycosidase activity having been shown by multiple groups and the activity can be increased by mutations to substitute for missing nucleophile or acid-base glutamate within the cavity (33). Furthermore, it has been demonstrated that sKLA binds sialyllactose of gangliosides to regulate membrane lipid raft formation and raft-dependent growth factor signaling (37). Moreover, molecular dynamic simulation reveals a slight movement of the β6α6 loop may “unblock” the entrance to allow potential substrate or ligand to gain access to the active site cavity (38). The large conformational changes of apo-sKLA we observed in 3DVA support the hypothesis. The factors controlling and influencing these conformational changes remain to be investigated.

One key difference between sKLA and sKLB is a metal binding site located between the sKLA KL1 and KL2 domains. This site likely serves to minimize interdomain flexibility and stabilize the KL2 domain as three of the residues are within it. Metal binding may favor FGF23 binding. Consistent with the notion, mutations to the four metal coordinating residues have been shown to disrupt FGF signaling (15). In the sKLA-FGF23-FGFR1c ternary complex crystal structure the bound metal was determined to be zinc. A search of the MetalPDB database for similar zinc containing sites in proteins with 3 Asp and 1 Cys as coordinating residues produced no hits. As zinc can be a common contaminant in many buffers used for purification, it may be worth considering if another metal could occupy the site in vivo and what functional significance it could have.

In conclusion, we reveal that sKLA exists in multiple conformational states and potentially multiple stoichiometry. These findings, together with the report by Zhong et al. that post-translational modifications impact function, provides additional support that sKLA is a FGF23-independent pleiotropic hormone and potentially with enzymatic activities. Much work is required to fully elucidate the biology and physiology of KLA in health and diseases.

## MATERIALS AND METHODS

### Size-exclusion chromatography

Recombinant sKLA (Glu34-Ser981) with a C-terminal 6His tag expressed in a NS0 murine myeloma cell line was purchased (R&D Systems) and loaded onto a Superdex 200 Increase 10/300 GL column (Cytiva) equilibrated in SEC buffer (20 mM HEPES pH 7.5, 100 mM ammonium sulfate, 500 mM NaCl, 5% glycerol, and with or without 10 mM dithiothreitol, DTT).

### Mass photometry

MP experiments were performed on a Refeyn TwoMP mass photometer (Refeyn Ltd, Oxford, UK). Microscope coverslips (24 mm x 50 mm, Thorlabs Inc.) were cleaned by serial rinsing with Milli-Q water and HPLC-grade isopropanol (Sigma Aldrich) followed by drying with a filtered air stream. Silicon gaskets (Grace Bio-Labs) to hold the sample drops were cleaned in the same procedure immediately prior to measurement. All MP measurements were performed at room temperature using buffer (20 mM HEPES, pH 7.5 and 150 mM NaCl). The instrument was calibrated using a protein standard mixture: β-amylase (Sigma-Aldrich, 56, 112 and 224 kDa), and thyroglobulin (Sigma-Aldrich, 670 kDa). Before each measurement, 15 µL of buffer was placed in the well to find focus. The focus position was searched and locked using the default droplet-dilution autofocus function after which 5 µL of protein samples (40 nM) was added and pipetted up and down to briefly mix before movie acquisition was promptly started. Movies were acquired for 60 s (3000 frames) using AcquireMP (version 2.3.0; Refeyn Ltd) using standard settings. All movies were processed and analyzed using DiscoverMP (version 2.3.0; Refeyn Ltd).

### Cryo-EM sample preparation and data acquisition

sKLA was vitrified on QuantiFoil R2/1 300 mesh copper grids in 25 mM HEPES pH 7.5, 300 mM NaCl, and 1 mM TCEP using a Vitrobot Mark IV (ThermoFisher Scientific). Grids were glow discharged for 60 s, –15 mA on a PELCO easiGlow (Ted Pella) system. Sample (3 mL) was applied to grids in the Vitrobot chamber (4 °C and 95% humidity) and blotted for three seconds before plunge-freezing in liquid ethane. Data were collected on a Titan Krios G3 microscope (300 kV) using SerialEM with a K3 direct electron detector (Gatan). A total of 8,676 movies were collected at a pixel size of 0.6507 Å/pixel (super-resolution mode) with a dose of ∼50 electrons/A^2^, exposure time of 1.099 seconds, 50 frames, and a defocus range of -1.0 to -2.5 mm.

The sKLA dimer was obtained while preparing a complex of sKLA-FGF23-FGFR1c by mixing in a molar ratio of 1:4:4 in 20 mM HEPES pH 7.5, 150 mM NaCl, and 1 mM DTT at 4C overnight. 0.2 mg/mL of mixture was vitrified on QuantiFoil R2/1 300 mesh copper grids using a Vitrobot Mark IV. Grids were glow discharged for 60 s, –15 mA on a PELCO easiGlow (Ted Pella) system. Sample (3 mL) was applied to grids in the Vitrobot chamber (4 °C and 95% humidity) and blotted for two seconds before plunge-freezing in liquid ethane. Data were collected on a Titan Krios G3 microscope (300 kV) using SerialEM with a K3 direct electron detector (Gatan). A total of 5,760 movies were collected at a pixel size of 0.53 Å/pixel (super-resolution mode) with a dose of ∼60 electrons/A^2^, exposure time of 2 seconds, 50 frames, and a defocus range of –0.8 to -2.5 mm.

### Cryo-EM image processing and 3D reconstruction

Movies were subject to patch motion correction and patch CTF estimation in cryoSPARC, initial 3D reconstructions were used to generate 2D templates for template-based picking on the full datasets. For sKLA monomer, particles (1,742,250) were extracted using a box size of 340 pixels (0.6507 Å/pixel) and cleaned with multiple rounds of 2D classification. Multiple classes were used for *ab initio* reconstruction followed by heterogeneous refinement to select 115,398 particles for non-uniform refinement in C1 symmetry. Processing was done initially in cryoSPARC v3.1 and finished in v3.3.1. Final maps were post-processed using DeepEMhancer. For sKLA dimer, particles (4,238,397) were extracted using a box size of 120 pixels (2.12 Å/pixel) and cleaned with multiple rounds of 2D classification. A smaller subset of particles (330,846) was selected for further 2D classification before selection and re-extraction of the final 30,219 particle stack with a final box size of 256 pixels (1.06 Å/pixel). These particles were used for *ab initio* reconstruction followed by non-uniform refinement in C1 symmetry. Processing was done in cryoSPARC v4.2.1.

### 3D variability analysis

To analyze variability present in sKLA particles, we used the 3D Variability Analysis (3DVA) job in cryoSPARC v3.3.1 (27). A stack of 844,899 particles from a non-uniform refinement and then the resulting particles and mask were used to compute 6 eigenvectors of the covariance matrix of the data distribution using 3DVA, with resolution filtered to 5 Å. Model series were generated using the 3DVA display job using 10 frames. To better analyze motion associated with the first two variability components, a series of models were generated corresponding to each volume series using *phenix.varref* (39). Supplemental movies were generated with Chimera and ChimeraX.

### Model building and refinement

An initial model for the sKLA consensus structure was generated by truncating regions from the sKLA-FGF23-FGFR1 complex structure (PDB ID 5W21). The model was initially docked into the density map using Fit in Map in Chimera (40). Manual model building was performed in Coot (41) and refinement using real-space refinement in Phenix (42). Figures were generated in Chimera and PyMOL (28). Software used for data processing, model building, and refinement, except for cryoSPARC, were curated by SBGrid (43).

### Accession number

Atomic coordinates and maps for apo-soluble klotho monomer and dimer have been deposited in the Protein Data Bank and Electron Microscopy Data Bank with the following accession code: 8TOH and EMD-41452 for the monomer; 8UF8 and EMD-42186 for the dimer.

## DATA AVAILABILITY

Requests for resources and reagents should be directed to and will be fulfilled by the lead contact, Chou-Long Huang. All data are available in the main text, methods, or supplemental information.

## ACKNOWLEDGMENTS

This work was primarily supported by the NIH National Institute of Diabetes and Digestive and Kidney Diseases grants DK100605 and DK109887 (to C.-L.H.). A portion of this research was supported by NIH grant U24GM129547 and performed at the PNCC at OHSU and accessed through EMSL (grid.436923.9), a DOE Office of Science User Facility sponsored by the Office of Biological and Environmental Research. We would like to thank Omar Davulcu for assistance with cryo-EM data collection at PNCC. Initial cryo-EM samples were prepared and imaged with assistance from Ed Brignole at the Automated Cryogenic Electron Microscopy Facility in MIT.nano on a Talos Arctica microscope, which was a gift from the Arnold and Mabel Beckman Foundation. Additional preliminary cryo-EM data was collected at the Iowa State University Cryo-EM facility on the Glacios 200kV TEM and Gatan K3 detector. We thank Dr. Puneet Juneja for his assistance in grid screening and initial data collection. We thank Dr. David Belnap at the University of Utah for his assistance in data collection setup for sKLA dimer. We would like to acknowledge the use of resources at the Carver College of Medicine’s Protein and Crystallography Facility at the University of Iowa.

## AUTHOR CONTRIBUTIONS

Conceptualization: C.L.H., L.G., N.S. Methodology: L.G., M.A., N.S., Z.X.

Investigation: L.G., M.A., N.S., Z.X.

Visualization: C.L.H., L.G., M.A., N.S., Z.X. Supervision: C.L.H.

Writing—original draft: M.A., N.S., Z.X.

Writing—review & editing: C.L.H., L.G., M.A., N.S., Z.X.

The file includes four Supplementary Movies:

**Supplementary Movie 1**: First variability component (C1) maps from 3DVA.

**Supplementary Movie 2**: Models fit to first variability component maps from 3DVA.

**Supplementary Movie 3**: Second variability component (C2) maps from 3DVA.

**Supplementary Movie 4**: Models fit to second variability component maps from 3DVA.

