## Supplemental Figures for "Conformational landscape of soluble α-klotho revealed by cryogenic electron microscopy"

**Conformational landscape of soluble a-klotho**

**revealed by cryogenic electron microscopy**

**Affiliations**

^#^Equal contribution

**Supplemental Figures**

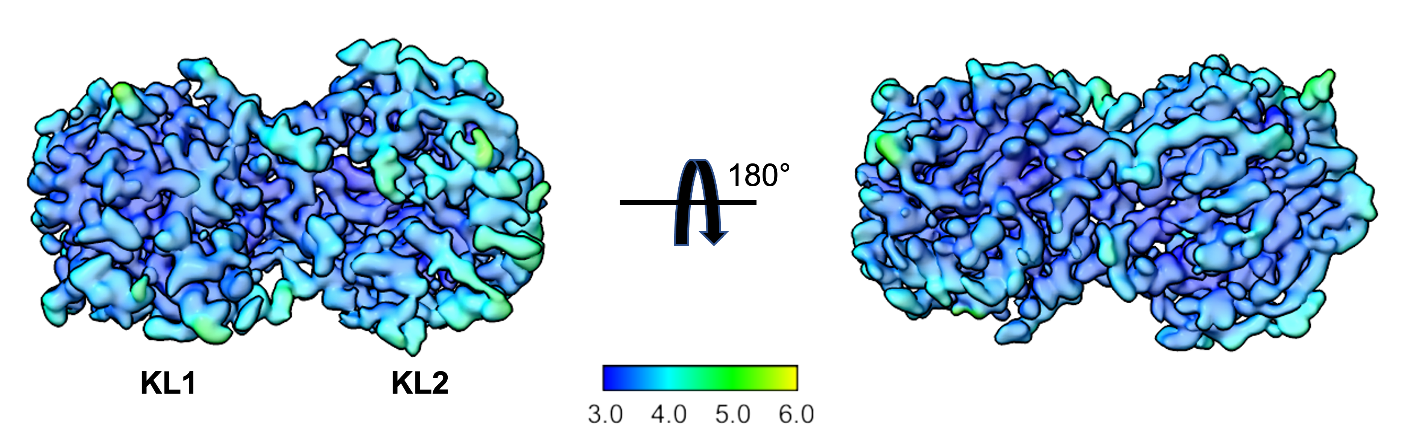

**Figure S1**. Local resolution cryo-EM map of the sKLA monomer. The local resolution ranges from 3.0 Å (inner regions of the KL1 domain) to ~5.0 Å near RBA loop surface regions in KL2 domain.

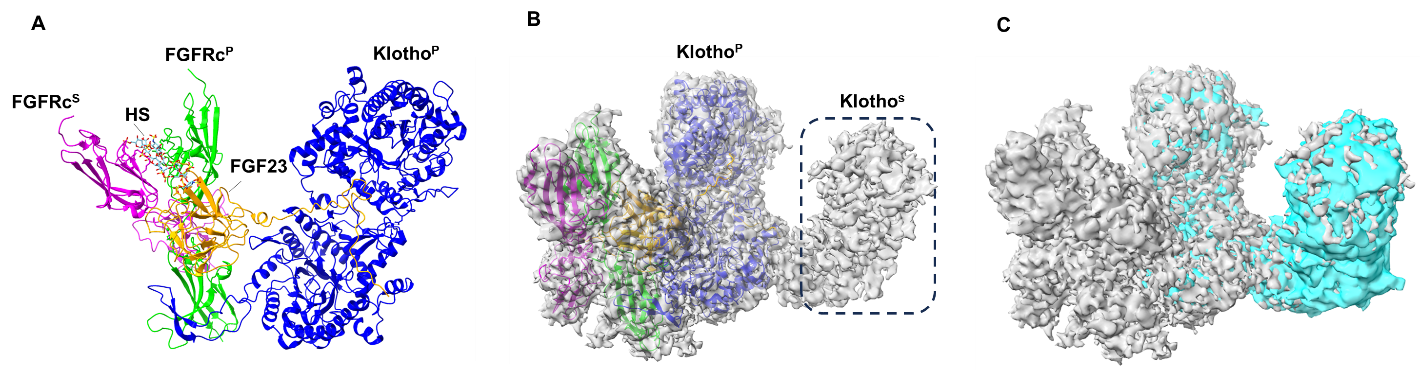

**Figure S2**. Cryo-EM map comparison of sKLA dimer in complex with or without FGF23-FGFR. Panel A and B are from reference (1) reporting asymmetric FGFR dimer formation. **(A)** Cartoon representation of FGF23-FGFR3c-αKlotho-HS (PDB: 7YSU) complex. For simplicity, the complex is shown in different colors, αKlotho^P^ (principal klotho, blue), FGF23 (orange), FGFRs^P^ (principal FGFR, lime) and FGFRs^S^ (secondary FGFR, magenta). **(B)** Cryo-EM maps of FGF23-FGFR3c-aKlotho-HS (grey, threshold level 0.34, EMDB-34082) with cartoon representation at 50% transparency. Extra weak density for a second klotho (αklotho^S^) is shown as dashed black box. **(C)** Density map fitting of apo-sKLA dimer from this study (cyan, threshold level 0.10) with Cryo-EM maps of FGF23-FGFR3c-aKlotho-HS.

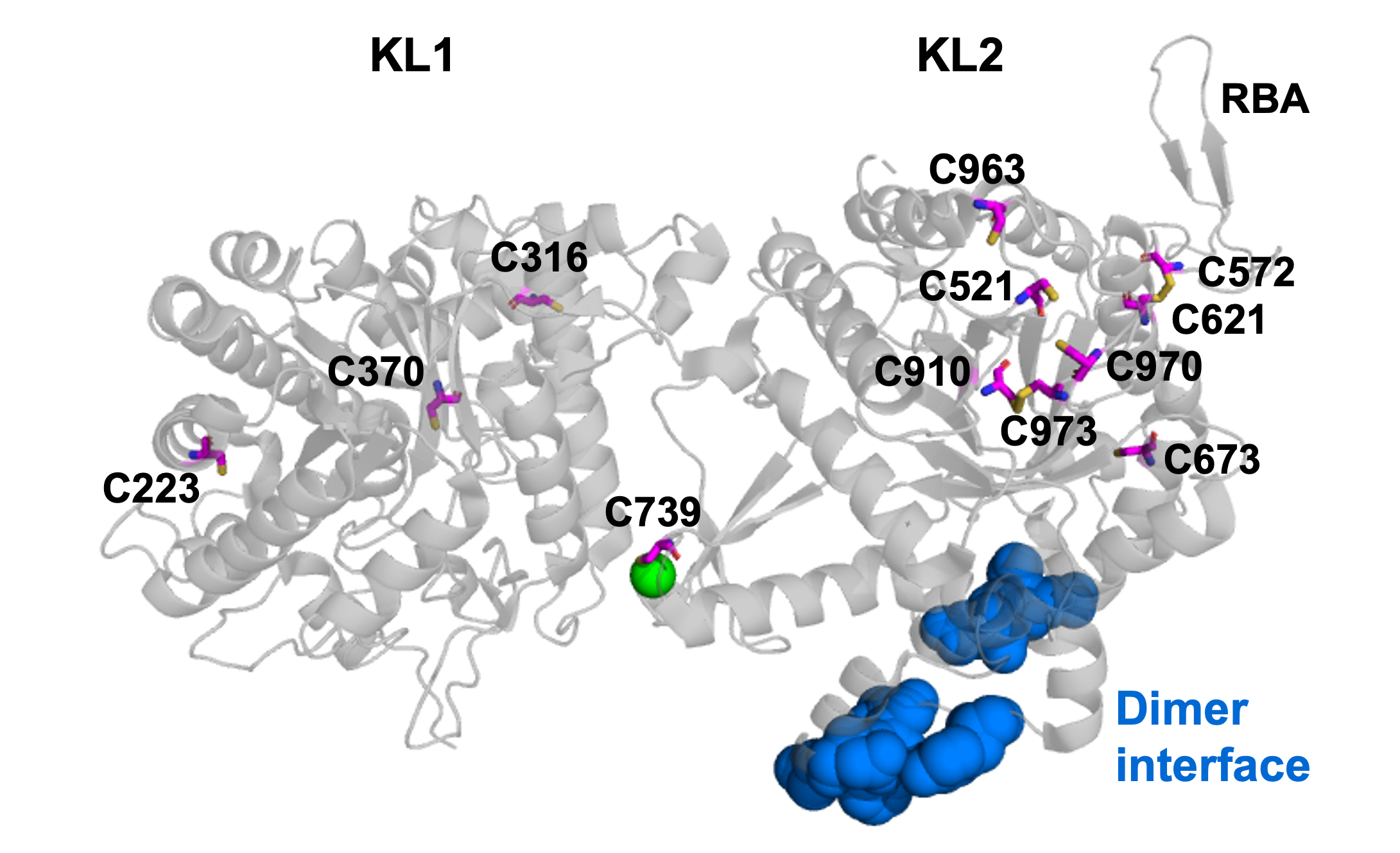

**Figure S3**. Cartoon representation of sKLA showing Cys residues (magenta sticks) and the dimerization interface (blue spheres). The green sphere is a zinc atom. Model is from PDB 5W21.

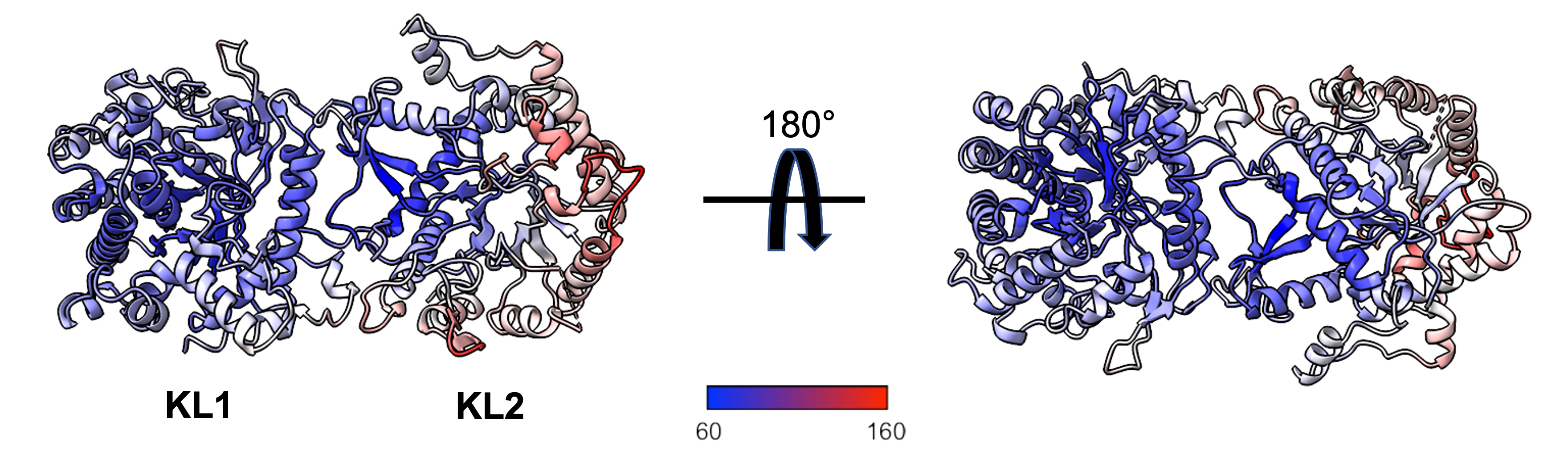

**Figure S4.** Debye–Waller (“B”) factors (quantifying vibrational motion of an atom) mapped onto the sKLA monomer cryo-EM structure. Blue regions represent lower B-factors and red indicates higher B-factors and regions with more movement. Scale bar (lowest to highest) ranges from 60-160 Å^2^.

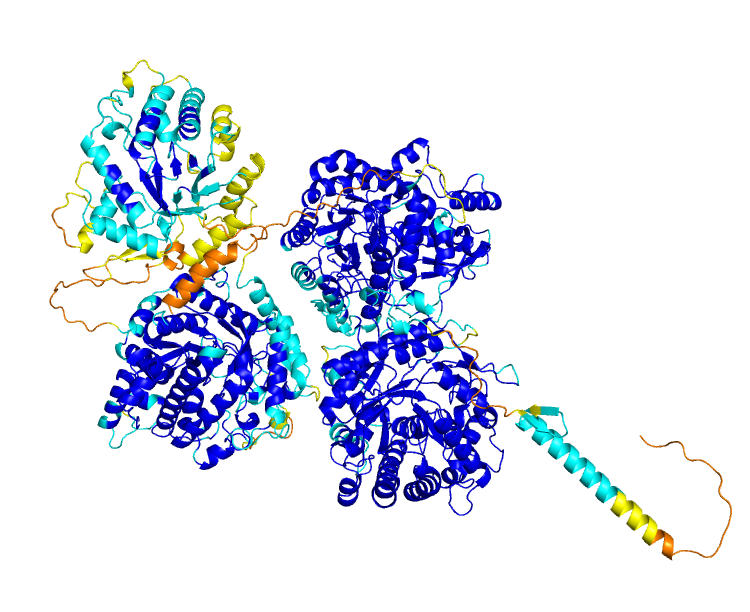

**Figure S5.** Alphafold database model for lactase-phlorizin hydrolase (UniProt: P09848) colored according to the confidence (pLDDT) score (Very high – blue, pLDDT > 90; High – cyan, 90 > pLDDT > 70; Low – yellow, 70 > pLDDT > 50; Very low – orange, pLDDT < 50).
